# *ZIP4* is required for normal progression of synapsis and for over 95% of crossovers in wheat meiosis

**DOI:** 10.1101/2023.03.17.532993

**Authors:** Tracie N. Draeger, María-Dolores Rey, Sadiye Hayta, Mark Smedley, Abdul Kader Alabdullah, Graham Moore, Azahara C. Martín

## Abstract

Tetraploid and hexaploid wheat have multiple genomes, with successful meiosis and preservation of fertility relying on synapsis and crossover only taking place between homologous chromosomes. In hexaploid wheat, the major meiotic gene *TaZIP4-B2* (*Ph1*) on chromosome 5B, promotes crossover between homologous chromosomes, whilst suppressing crossover between homeologous (related) chromosomes. Tetraploid wheat has three *ZIP4* copies: *TtZIP4-A1* on chromosome 3A, *TtZIP4-B1* on 3B and *TtZIP4-B2* on 5B. Previous studies showed that *ZIP4* mutations eliminate approximately 85% of crossovers, consistent with loss of the class I crossover pathway. Here, we show that disruption of two *ZIP4* gene copies in *Ttzip4-A1B1* double mutants, results in a 76-78% reduction in crossovers when compared to wild-type plants. Moreover, when all three copies are disrupted in *Ttzip4-A1B1B2* triple mutants, crossover is reduced by over 95%, suggesting that the *TtZIP4-B2* copy is also affecting class II crossovers. This implies that, in wheat, the class I and class II crossover pathways may be interlinked. When *ZIP4* duplicated and diverged from chromosome 3B on wheat polyploidization, the new 5B copy, *TaZIP4-B2*, may have acquired an additional function to stabilize both crossover pathways. In plants deficient in all three *ZIP4* copies, synapsis is delayed and does not complete, consistent with our previous studies in hexaploid wheat, when a similar delay in synapsis was observed in a 59.3Mb deletion mutant, *ph1b*, encompassing the *TaZIP4-B2* gene on chromosome 5B. These findings confirm the requirement of *ZIP4-B2* for efficient synapsis, and suggest that *TtZIP4* genes have a stronger effect on synapsis than previously described in Arabidopsis and rice. Thus, *ZIP4-B2* accounts for the two major phenotypes reported for *Ph1*, promotion of homologous synapsis and suppression of homeologous crossover.

**Key message:** In tetraploid wheat, *ZIP4* is required for efficient chromosome synapsis and for over 95% of crossovers, involving both the class I and class II crossover pathways.

**Author contribution statement:** TD grew and maintained the plants, made the crosses, carried out the KASP genotyping and sequencing, carried out the meiotic metaphase I studies and produced the corresponding figure, and wrote the manuscript. M-DR scored chromosome crossover, performed the statistical analysis and produced the graphs. AM selected the TILLING mutant, carried out the immunolocalization and FISH experiments and produced the immunolocalization figure; SH and MS developed the *Ttzip4-B2* CRISPR mutant in Kronos using RNA-guided Cas9 and produced the CRISPR *Ttzip4-B2* sequence figure; AKA designed the KASP primers; AM and GM provided the concept, provided thoughts and guidance, and revised and edited the manuscript.

**Conflict of interest:** The authors declare that they have no conflict of interest.

## Introduction

Bread wheat (*Triticum aestivum* L.) and pasta wheat (*Triticum turgidum* ssp. *durum*) are allopolyploids that arose by hybridization between different wheat progenitor species with related (homeologous) genomes. Bread wheat is a hexaploid (2n = 6x = 42), comprising three diploid sub-genomes (A, B and D), while pasta wheat is a tetraploid (2n = 4x = 28) comprising only two (A and B). The homeologous genomes possess a similar gene content and order. During early meiosis, maternal and paternal homologous chromosomes align as pairs, then become physically linked along their entire lengths in a process called synapsis, which is facilitated by the formation of the synaptonemal complex (SC) which assembles between them (Page and Hawley, 2004). Later, at metaphase I of meiosis, the 14 chromosomes of each sub-genome can be seen as seven bivalent pairs linked by chiasmata, the cytologically visible sites where chromosome crossover and recombination take place. Crossovers enable genetic information to be reciprocally exchanged between chromosomes to create new allelic combinations, with at least one crossover link between chromosome pairs (the obligate crossover) needed to ensure accurate chromosome segregation and balanced gametes in daughter cells (Zickler and Kleckner, 1999). Chromosome behaviour is strictly controlled, such that synapsis, crossover and recombination only occur between homologous chromosomes within each sub-genome and not between sub-genomes. Thus, polyploid wheat has evolved to behave cytologically as a diploid.

Several loci have been reported to help stabilise polyploid genomes during meiosis, including *PrBn* in *Brassica napus* and *Ph2* (*MSH7-3D*) in hexaploid wheat, both of which reduce homeologous crossover (Jenczewski et al., 2003; Serra et al., 2021). However, the strongest effect on the diploid behaviour of both tetraploid and hexaploid wheat has been previously ascribed to the *Ph1* (pairing homeologous 1) locus on chromosome 5B, which not only suppresses crossover between homeologous chromosomes (homeologs), but also promotes pairing and synapsis between homologous chromosomes (homologs) during early meiosis (Riley and Chapman, 1958; Sears and Okamoto, 1958; Wall et al., 1971; Martín et al., 2014 and 2017; Rey et al., 2017). Such mechanisms maintain genome stability and fertility, but for wheat breeding, the presence of *Ph1* was a barrier to the introgression of useful genes from wild relatives into modern wheat varieties, because crossover between the homeologous chromosomes of the two parents was suppressed. Two mutant lines with interstitial deletions of *Ph1* were subsequently used for breeding purposes: *ph1b* (Sears, 1977) in the hexaploid wheat variety ‘Chinese Spring’; and *ph1c* (Giorgi, 1978) in the tetraploid wheat variety ‘Cappelli’. In hybrids of these mutants with wild-relatives, high numbers of bivalent pairs were observed during meiosis, indicating that crossover was occurring between homeologous chromosomes. The deletion lines *ph1b* and *ph1c* were widely exploited for breeding purposes, however, after many generations of breeding, *ph1b* has accumulated extensive chromosomal rearrangements (Sánchez-Morán et al., 2001; Martín et al., 2018) resulting in reduced fertility and poor agronomic performance (Türkösi et al., 2022).

The *ph1b* deletion is now known to be 59.3 Mb, with 1,187 genes deleted (Martín et al., 2018). However, the effects of *Ph1* were further defined to a smaller region on chromosome 5BL (Roberts et al., 1999; Griffiths et al., 2006, Al-Kaff et al., 2008), now known to be 0.5 Mb (Martín et al., 2018). This 0.5 Mb region contains a cluster of *Cdk2-like* genes disrupted by a segment of heterochromatin incorporating a gene originally designated *Hyp3* (Griffiths et al., 2006; Al-Kaff et al., 2008), later reannotated as *TaZIP4-B2* (Martín et al., 2017; Rey et al., 2017). Although the exact mode of action is uncertain, *ZIP4* has a major role in meiosis: acting as a hub to facilitate interactions between components of the chromosome axis and proteins involved in the crossover process (De Muyt et al., 2018). In Arabidopsis (*Arabidopsis thaliana*) and rice (*Oryza sativa*) *ZIP4* is necessary for class I crossover formation (Chelysheva et al., 2007; Shen et al., 2012), whilst in budding yeast (*Saccharomyces cerevisiae*) it is required for synapsis as well as crossover (Tsubouchi et al., 2006). To determine whether *TaZIP4-B2* could be responsible for the *Ph1* phenotype, a *TaZIP4-B2* CRISPR mutant was analysed alongside the *ph1b* deletion mutant. In both mutants, around 56% of meiocytes exhibited abnormalities, with a correspondingly similar reduction in grain set (Alabdullah et al., 2021). This suggested that *TaZIP4-B2* promotes correct pairing of homologs. Moreover, when TILLING and CRISPR mutants of *TaZIP4-B2* were crossed with *Aegilops variabilis*, the hybrids showed levels of increased homeologous crossover similar to those previously reported in *ph1b-Ae. variabilis* hybrids (Rey et al., 2017 and 2018). Crossover was also similarly increased between related wheat chromosomes in *Tazip4-B2* and *ph1b* haploid mutants (Martín et al., 2021). This evidence is all consistent with *TaZIP4-B2* being the gene responsible for the effects of *Ph1*. However, although it is known that *Ph1* has a direct effect on synapsis (Martín et al., 2017), the role of *TaZIP4* in synapsis has not yet been established.

In addition to the *TaZIP4-B2* gene on chromosome 5B, hexaploid wheat carries a further three copies of *ZIP4* on group 3 chromosomes (3A, 3B and 3D). It is not yet known how these group 3 copies contribute to meiosis, though they are likely to promote the class I crossover pathway (Alabdullah et al., 2019). In contrast, tetraploid wheat has only three copies of *ZIP4*, on chromosomes 5B, 3A and 3B. Previous studies have shown that tetraploid (durum) wheat uses two pathways of meiotic recombination, the class I crossover pathway, accounting for ∼85% of meiotic crossovers, and the class II crossover pathway, responsible for the remaining ∼15% (Desjardins et al., 2020). In contrast to hexaploid wheat, little is known about chromosome synapsis and crossover in tetraploid wheat, with the role of the three tetraploid *ZIP4* copies yet to be elucidated. As well as being an important food crop, tetraploid wheat has fewer genomes, making it a simpler system with which to study meiosis, and allowing faster generation of complete null mutants. In the current study, we have used a TILLING population developed in the tetraploid wheat cultivar ‘Kronos’ (Krasileva et al., 2017), together with the CRISPR-Cas9 system, to generate a complete collection of single, double and triple *Ttzip4* mutants involving elimination/loss of function of one or more of the different *TtZIP4* copies. We have used a combination of cytogenetics and immunocytology to determine how disruption of these different *ZIP4* copies affects synapsis and crossover in tetraploid wheat.

## Materials and Methods

### Plant material and growth conditions

Tetraploid wheat *zip4* TILLING and CRISPR mutants were derived from the durum wheat cultivar ‘Kronos’ (*Triticum turgidum* 2n = 4x = 28; AABB), which was also used as a wild-type control. The TILLING line, Kronos3161 (Kr3161), containing an EMS-induced heterozygous mutation in *TtZIP4-A1*, was selected from the <u>Ensembl Plants</u> database (Bolser et al., 2016). The mutation is a splice donor variant (Variant ID: Kronos3161.chr3A.647481179), producing a premature stop codon just after the 3^rd^ intron. Kr3161 also has an EMS-induced homozygous missense mutation (Variant ID: Kronos3161.chr3B.672871007) in *TtZIP4-B1*. The tetraploid wheat (*T. turgidum)* cv.

Cappelli mutant *ph1c*, (Giorgi, 1978), which has a 59.3 Mb deletion of the *TtZIP4-B2* gene on chromosome 5B, was used as a *Ttzip4B2* single mutant. CRISPR-Cas9 technology was exploited to generate a single mutant for *TtZIP4-B2* with a single base pair deletion. Seeds were obtained from the Germplasm Resources Unit at the John Innes Centre: www.SeedStor.ac.uk.

*Ttzip4-B1* single mutants and *Ttzip4-A1B1* double mutants were generated by self-fertilising Kr3161 plants heterozygous for *TtZIP4-A1*. Kr3161 plants were also back-crossed with wild-type Kronos, and the Bc_1_M_1_ plants self-fertilized, to produce *TtZIP4-A1B1B2* control and *Ttzip4-A1* single mutant plants. Crosses were made between Kr3161 and Cappelli *ph1c* deletion mutants, to produce heterozygous M_1_ progeny that were self-fertilized to generate *Ttzip4-A1B2* double mutants, *Ttzip4-B1B2* double mutants and *Ttzip4-A1B1B2* triple mutants in the M_2_ generation. *ZIP4* genotypes were confirmed by KASP genotyping and Sanger sequencing. Plants were grown in a controlled environment room (CER) at 20 °C (day) and 15 °C (night) with a 16-hour photoperiod and 70% humidity. Following germination, Cappelli *ph1c* plants were given three-weeks of vernalization at 6-8 °C.

### Generation of *Ttzip4-B2* CRISPR mutants using RNA-Guided Cas9

CRISPR *Ttzip4-B2* mutants were generated in Kronos by the BRACT group at the John Innes Centre. Three single guide RNAs (sgRNA) specific to the hexaploid wheat *TaZIP4-B2* gene (Gene ID: *TraesCS5B02G255100*.*1)*, and previously reported in Rey et al., 2018 were used. The genomic DNA sequence of the target gene *TaZIP4-B2* from *T. aestivum* cv. ‘Fielder’ was compared by alignment to the *TtZIP4-B2* sequence of *T. turgidum* cv. ‘Kronos’ to confirm sgRNA validity. The specific *TtZIP4-B2* guides were: sgRNA 4: 5′ GATGAGCGACGCATCCTGCT 3′, sgRNA 11: 5′ GATGCGTCGCTCATCCTCCG 3′ and sgRNA 12: 5′ GAAGAAGGATGCGGCCTTGA 3′. Two binary vectors were prepared for wheat transformation using standard Golden Gate MoClo assembly (Werner et al., 2012). The Level 1 plasmids in positions 3 and 4, previously described in Rey et al., 2018, were reused for Level 2 assembly in this study. Each Level 1 plasmid contained a single guide RNA between the TaU6 promoter and the guide scaffold for *Streptococcus pyogenes* Cas9. Level 2 assembly was performed using the Level 2 acceptor pGoldenGreenGate-M (pGGG-M) (Addgene #165422) binary vector (Smedley et al., 2021). The Level 1 plasmids pL1P1OsActinP:*hpt*-int:35sT selection cassette (Addgene #165423), pL1P2OsUbiP:Cas9:NosT (Addgene #165424) and pL1P5ZmUbiP:GRF-GIF:NosT (Addgene #198046) and the sgRNA cassettes were assembled into pGGG-M along with end linker pELE-5 (Addgene #48020). The resulting plasmids were named pGGG-ZIP4-B2 Construct 1 (containing sgRNA 4 and 12) and pGGG-ZIP4-B2 Construct 2 (containing sgRNA 11 and 12). The two pGGG-ZIP4-B2 constructs were electroporated into *Agrobacterium tumefaciens* AGL1 (Lazo et al., 1991) competent cells and transformed into Kronos plants as described in Hayta et al., 2021. Transgene copy number was determined by Taqman qPCR and probe (Hayta et al., 2019), and used to calculate copy number according to Livak and Schmittgen (2001).

Primers used for screening of gene editing in the primary transgenics (T_0_) and subsequent generation (T_1_) are listed in Supplementary Table 1. T_0_ plants were screened by PCR amplification across the target regions followed by Sanger sequencing. Sequence chromatographs were visually analysed using Geneious Prime (Biomatters Ltd). Twenty-four T_1_ plants from 3 selected edited T_0_ lines were screened by PCR amplification of the target region followed by paired-end Illumina Next-Generation Sequencing (NGS) performed by Floodlight Genomics LLC (Knoxville, TN, USA). Sequencing reads were mapped to the Kronos *ZIP4-B2* reference sequence (<u>Grassroots Infrastructure</u>). Fastq files generated from bwa 0.7.17 were converted to bam files and further sorted and indexed with Samtools (1.10). The genome browser software Integrative Genomics Viewer (https://igv.org/app) was used to display NGS data for analysis. Eight homozygous edited plants were identified in the T_1_ generation. One plant, containing a single bp deletion (causing a frameshift in the amino acid sequence from codon 147 and a premature stop codon at 213), was chosen for further analysis.

### KASP genotyping of *ZIP4* wild type and mutant plants

Plants were grown to the 2-3 leaf stage, and DNA extracted from leaf material as in Draeger et al., 2020 (adapted from Pallotta et al., 2003). Extracted DNA was diluted with dH_2_0. Final DNA template concentrations were between 15-30 ng. KASP genotyping was performed using chromosome-specific primers designed from sequences from the Chinese Spring reference sequence assembly, IWGSC RefSeq v1.0 (International Wheat Genome Consortium, 2018). Primer sequences are shown in Supplementary Table 2. The allele-specific forward primers and common reverse primers were synthesized by Merck https://www.merckgroup.com/. Allele-specific primers were synthesized with standard FAM or VIC compatible tails at their 5’ ends (FAM tail: 5’ GAAGGTGACCAAGTTCATGCT 3’; VIC tail: 5’ GAAGGTCGGAGTCAACGGATT 3’).

### KASP reaction and PCR conditions for genotyping

The KASP reaction and its components were as recommended by LGC Genomics Ltd and described at https://www.biosearchtech.com/support/education/kasp-genotyping-reagents/how-does-kasp-work. Assays were set up as 5 μl reactions in a 384-well format and included 2.5 μl genomic DNA template (15-30 ng of DNA), 2.5 μl of KASP 2x Master Mix (LGC Genomics), and 0.07 μl primer mix. Primer mix consisted of 12 μl of each tailed primer (100 μM), 30 μl common primer (100 μM) and 46 μl dH_2_O. For most primers, PCR amplification was performed using the following program: Hotstart at 94 °C for 15 min, followed by ten touchdown cycles (94 °C for 20 s; touchdown from 65-57 °C for 1 min, decreasing by 0.8 °C per cycle) and then 30 cycles of amplification (94 °C for 20 s; 57 °C for 1 min). However, the *TtZIP4-A1* 3A-genome-specific primers have low melting temperatures (Tms), so for these primers the PCR program was adapted to: Hotstart at 94 °C for 15 min, followed by fifteen touchdown cycles (94 °C for 15 s; touchdown from 60-45 °C for 1 min, decreasing by 1 °C per cycle) and then 75 cycles of amplification (94 °C for 15 s; 45 °C for 30 s, 50 °C for 30 s). Fluorescent signals from PCR products were read in a PHERAstar microplate reader (BMG LABTECH Ltd.). If tight genotyping clusters were not obtained, additional rounds of 5 cycles were performed. Genotyping data was analysed using KlusterCaller software (LGC Genomics).

### Sequencing to distinguish *TtZIP4-A1 (3AA)* homozygotes and *TtZIP4-A1 (3Aa)* heterozygotes

Sequencing was carried out to distinguish between *TtZIP4-A1* homozygous wild type plants and heterozygotes, because these genotypes were not easily differentiated using KASP primers (*Ttzip4-A1* homozygous mutant genotypes always separated well using KASP). DNA samples were PCR amplified using the following primers: Forward primer: 5’ CCTACTGCTTCTTACGTTTGAC 3’; Reverse primer: 5’ CGTCCTCGTTGTTCTTCTG 3’.

The PCR program was: Hotstart at 94 °C for 10 min, followed by 35 cycles of amplification at 94 °C for 30 s; 61.5 °C for 1 min and 72 °C for 1 min; then a final extension of 72 °C for 10 min. After PCR amplification, products were separated using agarose gel electrophoresis, and DNA bands excised from the gel and cleaned using a Qiaquick Gel Extraction Kit (Qiagen Ltd., UK). Sanger sequencing was performed by Genewiz, UK (now Azenta Life Sciences, UK). Sequences were edited using the BioEdit Sequence Alignment Editor vs. 7.2.5.

### Meiotic metaphase I analysis

Anthers were sampled from immature spikes when plants had developed to between Zadoks growth stages 41 and 43 (Zadoks et al., 1974; Tottman, 1987), when meiosis is in progress. At this stage, the flag leaf had fully emerged, and excised spikes were between 3.5-5.5 cm in length (average 4.5 cm). Anthers were sampled from the first 5 tillers only. To identify anthers with meiocytes at metaphase I, one anther from each floret was stained with acetocarmine and squashed, and meiocytes were examined using a DM2000 light microscope (Leica Microsystems). The three anthers within a floret are synchronized in meiotic development, so when metaphase I chromosomes were identified in the first anther, the two remaining anthers from the same floret were prepared for cytological analysis by Feulgen staining with Schiff’s reagent as described by Draeger et al., (2020). Anthers were sampled from three plants of each genotype. For each plant, a minimum of 30 meiocytes were blind scored from the digital images. For each cell, the different meiotic chromosome configurations were counted. These were unpaired univalents (0 chiasmata), rod bivalents (1-2 chiasma), ring bivalents (2-3 chiasmata), trivalents (2–3 chiasmata) and tetravalents (3 chiasmata). Chiasma frequency per meiocyte was calculated separately using two different methods, to give single chiasmata scores representing the minimum number of chiasmata per cell and double chiasmata scores representing the maximum. Statistical analyses were performed using STATISTIX 10.0 software (Analytical Software, Tallahassee, FL, USA).

All lines were analysed by the Kruskal-Wallis test (nonparametric one-way analysis of variance). Means were separated using Dunn’s test with a probability level of 0.05. Column charts were plotted using Microsoft Excel (2016).

### Immunolocalization of ASY1 and ZYP1 and FISH labeling of telomeres

Immunolocalization of antibodies against the meiotic proteins ASY1 and ZYP1 was combined with labeling of telomeres by fluorescence *in situ* hybridization (FISH), to follow the progression of synapsis. Anthers at the desired stages of meiosis were collected from Kr3161 *TtZIP4-A1B1B2* (wild-type control), *Ttzip4-A1B1* (double mutant) and *Ttzip4-A1B1B2* (triple mutant) plants at selected stages of meiosis, as described above for meiotic metaphase I, except that for immunolocalization combined with FISH, anthers were fixed in paraformaldehyde 4%/0.5% Triton™ X-100 for 15 min and processed immediately. Anthers were tapped in 1xPBS to release the meiocytes, and 20μl of this suspension were transferred onto a Polysine slide (poly-L-lysine coated slide) and left to air dry at room temperature for around 15 min. To preserve the 3D structure of the cells, no pressure was applied on the meiocyte suspension.

Antibodies do not always tolerate the aggressive procedures carried out during FISH, so immunolocalization was conducted first, followed by a gentle FISH procedure as described below. Slides were incubated in a detergent solution for 20 min (1xPBS, 0.5% Triton, 1mM EDTA) and blocked in 3% bovine serum albumin (in 1xPBS, 0.1% Tween 20, 1mM EDTA) for 30 min, before being incubated in the primary antibody solution for 1 h at room temperature, followed by 48 h incubation at 4 ºC. The primary antibody solution consisted of Anti-TaASY1 (Boden et al., 2009) raised in rabbit, used at a dilution of 1:200 (in 1xPBS), and anti-HvZYP1 (Colas et al., 2016) raised in rat and used at a dilution of 1:200 (in 1xPBS). Slides were kept at room temperature for 1 h, washed in 1xPBS and incubated with the secondary antibody for 1 h at 37 ºC. Anti-rabbit Alexa Fluor® 488 (Thermo Fisher Scientific, #A-11008) and anti-rat Alexa Fluor® 568 (Thermo Fisher Scientific, #A-11077) diluted in 1xPBS were used as secondary antibodies. Following immunolocalization, slides were washed in 1x PBS for 15 min and re-fixed in paraformaldehyde 4% for 1 h. FISH was carried out as previously described (Martín et al., 2017), but denaturation of the slides was reduced to 7 min at 70 ºC. A telomere repeat sequence (TRS) probe was amplified by PCR as described previously (Cox et al., 1993) and labeled using the biotin-nick translation mix (Roche Applied Science, # 11745824910), according to the manufacturer’s instructions. The biotin-labeled probe was detected with streptavidin, Alexa Fluor™ 660 conjugate (Invitrogen, #S21377). Slides were counterstained with DAPI (1μg/mL), mounted in Prolong Diamond antifade reagent (Thermo Fisher Scientific, #P36961) and left to cure for 2 or 3 days (to reach an optimum 1.47 refractive index) before images were collected.

### Image acquisition and analysis

Images of the metaphase I chromosomes were captured using a DM2000 microscope equipped with a DFC450 camera and controlled by LAS v4.4 system software (Leica Microsystems). For each cell, images were captured in up to 8 different focal planes to aid scoring.

Prophase I meiocytes labeled by FISH and immunofluorescence were optically sectioned using a DM5500B microscope (Leica Microsystems), equipped with a Hamamatsu ORCA-FLASH4.0 camera and controlled by Leica LAS-X software v2.0. Z-stack images of the meiocytes were processed using the deconvolution module of the Leica LAS-X software package. Images were further processed using Fiji (an implementation of ImageJ), a public domain program by W. Rasband available from http://rsb.info.nih.gov/ij/ (Schneider et al., 2012).

## Results

### Cytogenetic analysis of *zip4* single mutants at meiotic metaphase I

Anthers were sampled from three plants of each genotype, and a minimum of 30 meiocytes scored for each plant (at least 100 cells scored per genotype). Meiotic metaphase I chromosomes were blind scored for the numbers of univalents, ring and rod bivalents, trivalents and tetravalents, and for single and double chiasmata. Representative images of metaphase I chromosomes for each genotype and examples of the scored structures are shown in Figure 1. Statistical comparisons of the means are shown in Table 1 and Supplementary Table 3. Results are represented graphically in Figures 2 and Supplementary Figure 1.

**Figure 1.**
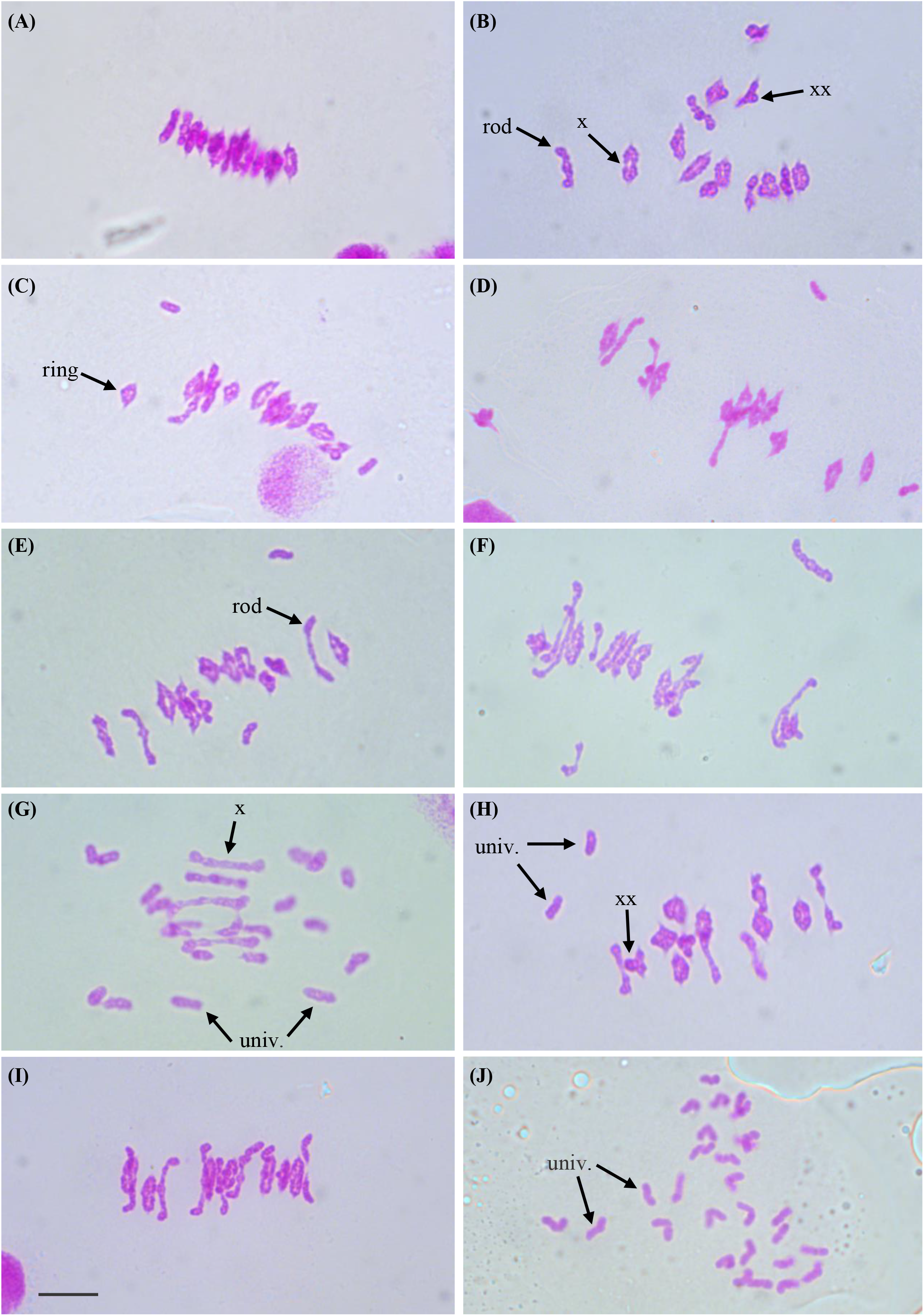
Representative images of chromosomes at metaphase I of meiosis from meiocytes of Kronos plants with differing *TtZIP4* genotypes (A) Wild-type Kronos; (B) Kr3161 *TtZIP4-A1B1B2*; (C) *Ttzip4-A1* single mutant; (D) *Ttzip4-B1* single mutant; (E) *Ttzip4-B2* single mutant (*ph1c*); (F) CRISPR *Ttzip4-B2* single mutant; (G) *Ttzip4-A1B1* double mutant; (H) *Ttzip4-A1B2* double mutant; (I) *Ttzip4-B1B2* double mutant; (J) *Ttzip4-A1B1B2* triple mutant. Examples of univalent chromosomes (univ.), rod bivalents (rod), ring bivalents (ring) bivalents (ring), single chiasmata (X) and double chiasmata (XX) are indicated with arrows. Note the greatly increased univalence in the *Ttzip4-A1B1* double mutant (G) and complete univalence in the *Ttzip4-A1B1B2* triple mutant (J). Scale bars, 10 μm.

**Table 1.** Genotypic effects on meiotic metaphase I chromosomes of *Ttzip4* single, double and triple mutants compared with those of Kr3161 *TtZIP4-A1B1B2* control plants. The mean numbers of univalents, rod and ring bivalents, trivalents and tetravalents were scored along with chiasma frequency scored as single and double chiasmata. Standard error (SE) values are shown. The range is given in brackets. P-values < 0.05 indicate significant differences. Superscript letters a-e indicate where the significant differences lie. For scores with the same letter, the difference between the means is not statistically significant. If the scores have different letters, they are significantly different. Note particularly the high numbers of univalents in the *Ttzip4-A1B1* double mutants and almost total univalence in the *Ttzip4-A1B1B2* triple mutants, and the corresponding low levels of chiasma frequency.

| Genotype | No. of | Univalents | Rod bivalents | Ring bivalents | Trivalents | Tetravalents | Single chiasmata | Double chiasmata |
| --- | --- | --- | --- | --- | --- | --- | --- | --- |

| | cells<br>scored | Mean ± SE<br>(Range) | Mean ± SE<br>(Range) | Mean ± SE<br>(Range) | Mean ± SE<br>(Range) | Mean ± SE<br>(Range) | Mean ± SE<br>(Range) | Mean ± SE<br>(Range) |
| --- | --- | --- | --- | --- | --- | --- | --- | --- |
| *ZIP4-A1B1B2* | 168 | 0.31 ± 0.06<sup>e</sup><br>(0-2) | 1.74 ± 0.10<sup>d</sup><br>(0-6) | 12.10 ± 0.10<sup>a</sup><br>(8-14) | 0.00 ± 0.00<sup>a</sup><br>(-) | 0.00 ± 0.00<sup>a</sup><br>(-) | 25.94 ± 0.12<sup>a</sup><br>(21-28) | 28.07 ± 0.12<sup>a</sup><br>(23-31) |
| *zip4-A1* | 127 | 0.57 ± 0.09<sup>de</sup><br>(0-6) | 1.76 ± 0.11<sup>d</sup><br>(0-6) | 11.95 ± 0.11<sup>a</sup><br>(7-14) | 0.00 ± 0.00<sup>a</sup><br>(-) | 0.00 ± 0.00<sup>a</sup><br>(-) | 25.70 ± 0.14<sup>a</sup><br>(20-28) | 27.42 ± 0.14<sup>a</sup><br>(22-30) |
| *zip4-B1* | 128 | 1.46 ± 0.14<sup>bc</sup><br>(0-9) | 3.16 ± 0.13<sup>bc</sup><br>(0-9) | 10.13 ± 0.15<sup>bc</sup><br>(5-14) | 0.00 ± 0.00<sup>a</sup><br>(-) | 0.00 ± 0.00<sup>a</sup><br>(-) | 23.43 ± 0.20<sup>bc</sup><br>(17-28) | 25.30 ± 0.17<sup>b</sup><br>(20-29) |
| *zip4-B2*<br>(*ph1c*) | 144 | 1.96 ± 0.12<sup>bc</sup><br>(0-6) | 3.89 ± 0.13<sup>ab</sup><br>(1-9) | 9.13 ± 0.14<sup>cd</sup><br>(5-13) | 0.00 ± 0.00<sup>a</sup><br>(-) | 0.00 ± 0.00<sup>a</sup><br>(-) | 22.17 ± 0.17<sup>cd</sup><br>(18-27) | 25.04 ± 0.20<sup>b</sup><br>(20-31) |
| *zip4-B2*<br>CRISPR | 157 | 2.07 ± 0.13<sup>bc</sup><br>(0-8) | 4.17 ± 0.13<sup>a</sup><br>(0-9) | 8.76 ± 0.14<sup>d</sup><br>(4-13) | 0.02 ± 0.01<sup>a</sup><br>(0-1) | 0.01 ± 0.01<sup>a</sup><br>(0-1) | 21.75 ± 0.18<sup>d</sup><br>(15-26) | 22.68 ± 0.19<sup>c</sup><br>(16-27) |
| *zip4-A1B1* | 164 | 17.77 ± 0.33<sup>a</sup><br>(8-26) | 4.48 ± 0.15<sup>a</sup><br>(0-8) | 0.63 ± 0.06<sup>e</sup><br>(0-4) | 0.00 ± 0.00<sup>a</sup><br>(-) | 0.00 ± 0.00<sup>a</sup><br>(-) | 5.75 ± 0.20<sup>e</sup><br>(1-12) | 6.64 ± 0.24<sup>d</sup><br>(0-14) |
| *zip4-A1B2* | 108 | 1.22 ± 0.12<sup>cd</sup><br>(0-4) | 2.83 ± 0.15<sup>c</sup><br>(0-7) | 10.56 ± 0.17<sup>b</sup><br>(6-14) | 0.00 ± 0.00<sup>a</sup><br>(-) | 0.00 ± 0.00<sup>a</sup><br>(-) | 23.94 ± 0.20<sup>b</sup><br>(19-28) | 25.09 ± 0.22<sup>b</sup><br>(19-29) |
| *zip4-B1B2* | 156 | 2.28 ± 0.14<sup>b</sup><br>(0-8) | 3.96 ± 0.12<sup>a</sup><br>(1-8) | 8.88 ± 0.13<sup>d</sup><br>(5-13) | 0.01 ± 0.01<sup>a</sup><br>(0-1) | 0.01 ± 0.01<sup>a</sup><br>(0-1) | 21.75 ± 0.17<sup>d</sup><br>(17-27) | 23.83 ± 0.30<sup>c</sup><br>(17-40) |
| *zip4-A1B1B2* | 128 | 25.88 ± 0.20<sup>a</sup><br>(18-28) | 1.04 ± 0.10<sup>d</sup><br>(0-5) | 0.02 ± 0.01<sup>e</sup><br>(0-1) | 0.00 ± 0.00<sup>a</sup><br>(-) | 0.00 ± 0.00<sup>a</sup><br>(-) | 1.08 ± 0.10<sup>e</sup><br>(0-5) | 1.20 ± 0.13<sup>d</sup><br>(0-8) |
| *p-value* |  | < 0.0001 | < 0.0001 | < 0.0001 | 0.0361 | 0.6277 | < 0.0001 | < 0.0001 |

**Figure 2.**
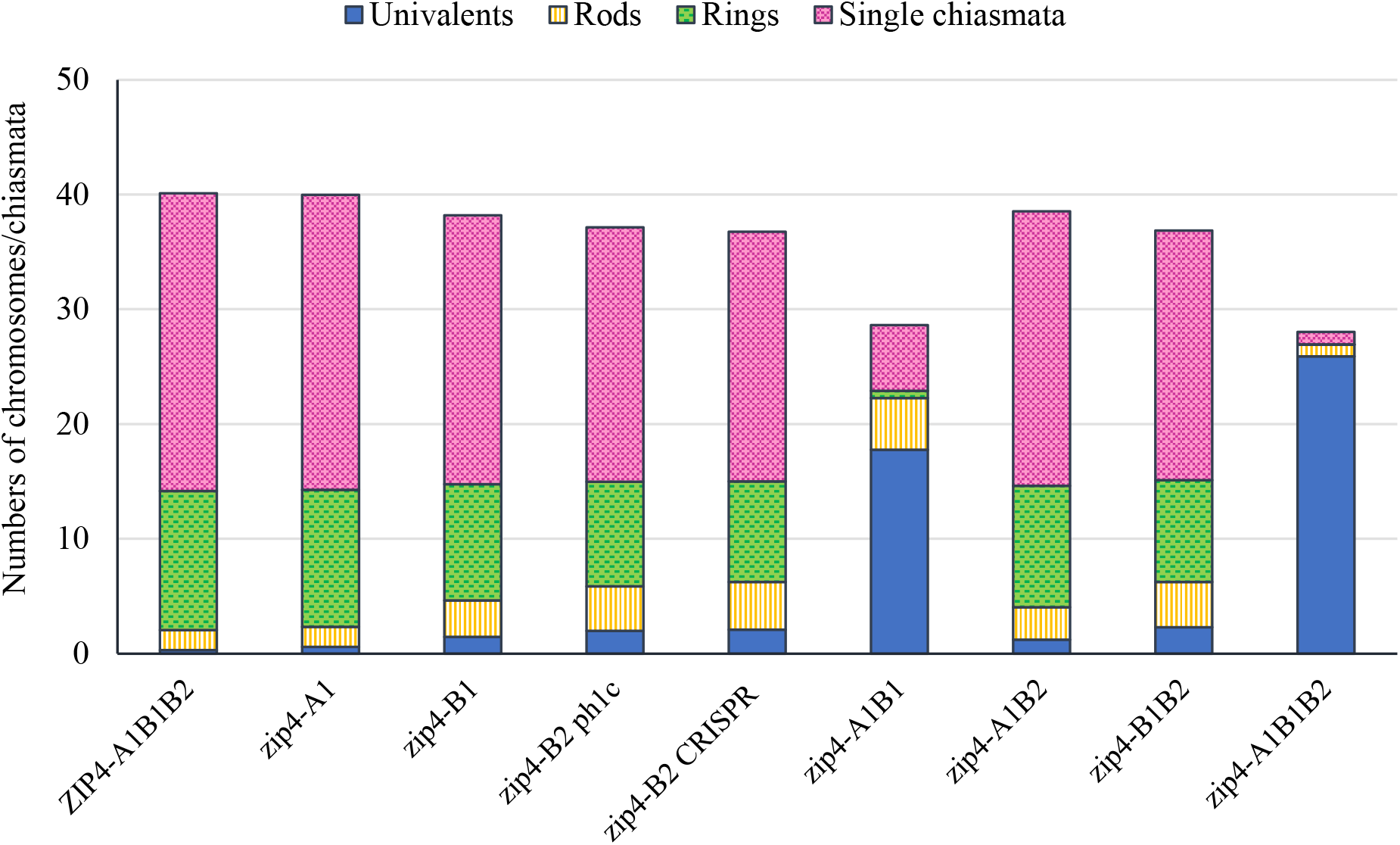
Column chart showing genotypic effects on meiotic metaphase I chromosomes of tetraploid wheat *Ttzip4* mutants compared with Kr3161 wild type control plants (*ZIP4-A1B1B2)*. Numbers of univalents, rod and ring bivalents and single chiasmata are shown. Numbers of multivalents and double chiasmata are not shown. Note greatly increased numbers of univalents (∼18 out of 28 chromosomes) and low chiasma frequency in *zip4-A1B1* double mutants, and virtually complete univalence and almost no chiasmata in the *zip4-A1B1B2* triple mutants.

First, chromosome scores from wild-type Kronos plants were compared with those of the Kr3161 *TtZIP4-A1B1B2* (wild type with mutant background) plants (Supplementary Table 3; Supplementsary Figure 1). Both genotypes had an average of around 12 ring bivalents and two rod bivalents per meiocyte (Figures 1A and B). Univalent chromosomes occurred only occasionally. The only significant difference between these lines was a slight decrease in single and double chiasma frequency in the Kr3161 control, but the means were similar: 26.25 and 25.94 for single chiasmata in wild-type Kronos and the Kr3161 control respectively, and 28.76 and 28.07 for double chiasmata.

Based on these analyses, we compared the *Ttzip4* mutant genotypes with the Kr3161 *ZIP4-A1B1B2* control lines (Table 1; Figure 2), given that most of the mutant genotypes should be more similar to this line than to wild-type Kronos plants in terms of their background mutations. Statistical analysis identified significant differences between genotypes for all chromosome configurations, except for trivalents and tetravalents, which were rare and confined to the CRISPR *Ttzip4-B2* and *Ttzip4-B1B2* mutants. There were no significant differences between *Ttzip4-A1* single mutants and wild type plants for any chromosome configurations, although univalent numbers ranged from 0-2 per meiocyte in wild type and 0-6 in *Ttzip4-A1* single mutants. *Ttzip4-B1* single mutants had significantly higher numbers of univalents and rod bivalents, and significantly fewer ring bivalents and chiasmata compared to control plants. Chiasma frequency was reduced by around 10%.

Of the single mutants, the largest effect was seen in *Ttzip4-B2*. In the CRISPR *Ttzip4-B2* mutant, for example, significant differences were seen for all chromosome conformations except for multivalents. Mean numbers of ring bivalents reduced from around twelve (12.10) in the Kr3161 control to nine (8.76) in the CRISPR mutant, with rods increasing from around two (1.74) to around four (4.17), and univalents increasing from less than one (0.31) to around two (2.07). In the CRISPR mutant, single (21.75) and double (22.68) chiasma frequency was also significantly lower when compared with control plants (25.94 single, 28.07 double), a decrease of 16-19%. Although the *Ttzip4-B2* CRISPR and *ph1c* mutants have different backgrounds (Kronos and Cappelli respectively), and the CRISPR mutant has lower levels of background mutations, the only significant difference between these two lines was in double chiasma frequency, where the mean was 25.04 for the *ph1c* mutant and 22.68 for the CRISPR mutant, a decrease of 14% in comparison to the Kr3161 wild type. A few multivalent chromosomes were observed in the CRISPR mutant and none in the *ph1c* mutant, but this difference was not statistically significant. Only two trivalents and one tetravalent chromosome were observed in the CRISPR mutant, out of 157 meiocytes scored.

### Reduced chiasma frequency and sterility in *zip4* double and triple mutants

In the *Ttzip4-A1B2* double mutants, numbers of univalents, bivalents and chiasma frequencies were similar to those seen in the *Ttzip4-B1* single mutants, with a reduction in chiasma frequency of 8-11%. *Ttzip4-B1B2* double mutants had similar scores to CRISPR *Ttzip4-B2* single mutants, with significantly more univalents and rod bivalents, and significantly fewer ring bivalents and chiasmata than *Ttzip4-A1B2* double mutants. However, the most striking differences between genotypes were seen in the *Ttzip4-A1B1* double mutants, which showed a 76-78% reduction in chiasma frequency and an average of around 18 univalents (compared to less than one in Kr3161 control plants), and in the *Ttzip4-A1B1B2* triple mutants, which showed a 96% reduction in chiasma frequency and an average of around 26 univalents, with 50 meiocytes out of 128 (almost 40%) showing complete univalence. Numbers of ring bivalents for both these genotypes fell from around twelve in the Kr3161 control to less than one. *Ttzip4-A1B1* double mutants had the highest number of rod bivalents (4.48) of all the genotypes, but rods were hardly ever observed in the triple mutant (1.04) because most chromosomes were univalent. Phenotypes of these two mutants were clearly distinguishable from those of wild-type Kronos (Figure 1A) and the Kr3161 control (Figure 1B) in the cell images. In the *Ttzip4-A1B1* double mutant, any rod bivalents present generally aligned on the metaphase I plate, with univalents spread out on either side, orientated to different poles of the nucleus (Figure 1G). This arrangement was also seen in the triple mutant when rods were present, but when meiocytes contained univalents alone, these were often dispersed across the cell, as in Figure 1J. Asynchrony and mis-segregation were evident in the triple mutant during stages other than metaphase I. All of the *Ttzip4-A1B1* double mutants and the *Ttzip4-A1B1B2* triple mutants were completely sterile.

### Delayed and incomplete synapsis in *Ttzip4-A1B1B2* triple mutants

During early meiosis, telomeres cluster as a bouquet at one pole of the nucleus, and homologous chromosomes pair intimately and synapse from these telomere regions. Previously we showed that in the *ph1b* mutant (which lacks *TaZIP4-B2*), progression of synapsis is slower than in the wild type, which allows some homeologous synapsis to take place (Martín et al., 2017). Therefore, we investigated the effect of eliminating different *TtZIP4* copies on the dynamics of synapsis. To track synapsis, we combined FISH labeling of telomeres with immunolocalization of the meiotic proteins ASY1 and ZYP1. ASY1 localises to regions of chromatin that associate with the axial/lateral elements of meiotic chromosomes, and is loaded before synapsis begins (Armstrong et al., 2002; Boden et al., 2009). In wheat, ASY1 is observed in regions that are not synapsed (Martín et al., 2014). ZYP1 is part of the central element of the synaptonemal complex (SC), a component of the transverse filaments that are installed between lateral elements as the SC assembles (Higgins et al., 2005; Khoo et al., 2012), and it is only present in chromosome regions that are synapsed. ZYP1 signal only lengthens into regions of chromatin after the ASY1 signal has unloaded.

In this study, ASY1, ZYP1 and telomere dynamics were monitored throughout meiotic prophase I in the *Ttzip4-A1B1* double mutant, the *Ttzip4-A1B1B2* triple mutant and the wild-type control. Figure 3A shows telomeres (labeled in red) clustered together at one pole of the nucleus, indicating that the telomere bouquet has formed in all wild-type and mutant meiocytes, with no difference observed between genotypes. In wild-type wheat at this early stage, ASY1 labeling (in magenta) can be seen across most of the nucleus, while ZYP1 tracks (in green) show the typical ZYP1 polarization (Martín et al., 2017), indicating that ZYP1 is polymerising from the nuclear pole containing the telomere bouquet. In the double mutant, where the group 3 *ZIP4* copies have been eliminated, ASY1 and ZYP1 loading is similar to that observed in the wild-type control, with the classical ZYP1 polarization starting from the telomere bouquet. However, in the absence of all three *TtZIP4* copies in the triple mutant, synapsis initiation is clearly delayed: while ASY has loaded normally and can be seen throughout the nucleus, ZYP1 does not show the usual polymerization starting from the telomere bouquet, and only short stretches of ZYP1 are observed dispersed throughout the nucleus (Figure 3A).

**Figure 3.**
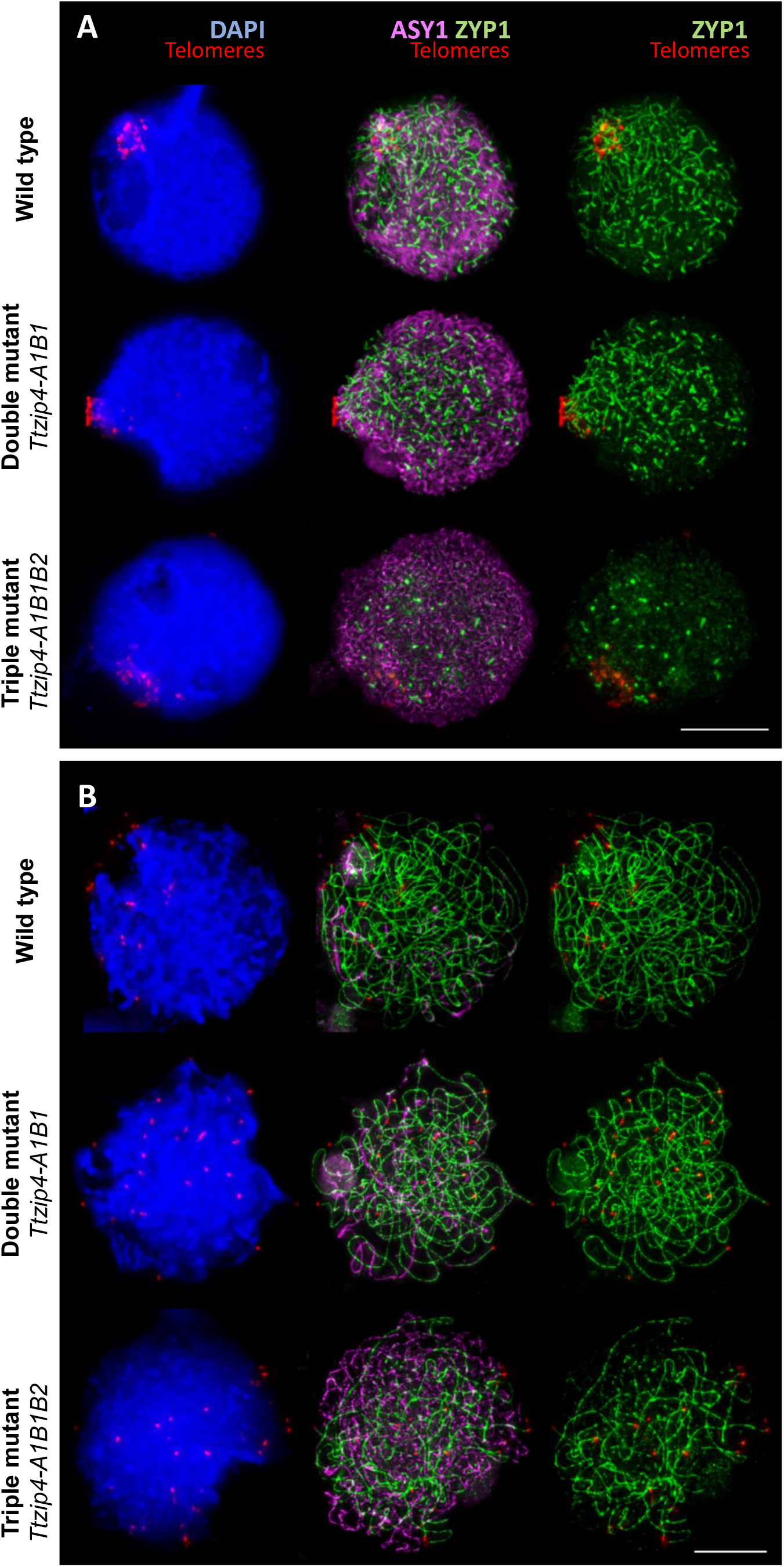
Immunolocalization of the meiotic proteins ASY1 (magenta) and ZYP1 (green) combined with the telomeres (red) labeled by FISH in tetraploid wheat with different copies of *TtZIP4* (*TtZIP4-A1B1B2* (wild-type control), *zip4A1B1* (double mutant) and *zip4-A1B1B2* (triple mutant). DNA DAPI staining in blue. Scale bar represents 10 μm. (**A**) Synapsis during the early telomere bouquet stage. Initiation of synapsis is observed at the telomere bouquet in the wild type and double mutant showing the typical ZYP1 polarization. However, in the triple mutant, in the absence of all *ZIP4* copies, synapsis initiation is mostly delayed and only short stretches of ZYP1 are observed dispersed throughout the nucleus. (**B**) Progression of synapsis after bouquet dispersal. Synapsis in the control and double mutant is almost completed at this stage, while synapsis is far from completion in the absence of all *ZIP4* copies, as illustrated by the large amount of ASY1 labeling still visible.

To assess whether there was any difference in synapsis between wild-type and the double mutant, and whether synapsis in the triple mutant had recovered to normal levels later in prophase I, we analysed the progression of synapsis, from initiation of telomere bouquet formation to complete telomere bouquet dispersal. Interestingly, we did not observe any difference in synapsis between wild-type and the double mutant (Figure 3B). After bouquet dispersal, synapsis was almost completed in both wild-type and double mutant, as can be observed by the very small amount of ASY1 labeling still visible at pachytene, representing the small amount of chromatin not yet synapsed (Figure 3B). In the experiments described here, only snapshots of a very dynamic process were taken, so small changes in synapsis dynamics between wild type and double mutants cannot be ruled out; however, no major differences were identified in terms of the extent of synapsis or its dynamics. In contrast, in the triple mutant, where all *ZIP4* copies were eliminated, there was a clear delay in synapsis, as illustrated by the large amount of ASY1 labeling still visible and the smaller amount of ZYP1 labeling compared to the wild type or double mutant. This indicates that synapsis has been compromised and is not completed (Figure 3B). No meiocyte was observed with synapsis even close to completion.

## Discussion

### *ZIP4* is required for over 95% of wheat meiotic crossovers

In most eukaryotes, including plants, two main pathways of meiotic crossover formation exist (Higgins et al., 2004; Mercier et al., 2005). The class I pathway produces crossovers subject to ‘interference’, which prevents crossovers from occurring close to each other (Jones and Franklin, 2006; Berchowitz and Copenhaver 2010), whereas in the class II pathway, there is no interference between crossovers, so they are randomly distributed along chromosomes (de los Santos et al., 2003; Osman et al., 2003; Higgins et al., 2008). The class I pathway accounts for the majority of crossovers in most examined species, and this pathway that ensures that each chromosome pair receives at least one crossover (the ‘obligate’ crossover), which promotes correct chromosome segregation (Jones and Franklin, 2006). In budding yeast, Arabidopsis and wheat, mutant studies have shown that around 85% of crossovers are formed via the class I pathway and around 15% via the class II pathway (de los Santos et al., 2003; Börner et al., 2004; Tsubouchi et al., 2006; Chelysheva et al., 2007; Desjardins et al., 2020).

Formation of class I crossovers is controlled by a group of conserved, meiosis-specific proteins called ZMMs, named after the budding yeast proteins ZIP1-4, MSH4-5 and MER3 (Börner et al., 2004; Reviewed Mercier et al., 2015). Removing ZMM proteins can result in a drastically altered number of class I crossovers or their complete absence (Pyatnitskaya et al., 2019). Disruption of *ZIP4* genes in diploid species, where genes are often present as single copies, usually results in sterility, as elimination of homologous crossovers leads to incorrect segregation. For example, in Arabidopsis and rice, mutations in *ZIP4* eliminate around 85% and 70% of crossovers respectively, and *zip4* mutants are mostly sterile (Chelysheva, 2007; Shen et al., 2012). In contrast, polyploids often have two or more gene copies that perform the same function, and inactivation of one of these copies may have little or no effect on the phenotype. For example, in tetraploid wheat, single mutants of the meiosis-specific gene *MSH4* are fully fertile, whereas *Ttmsh4ab* double mutants are sterile (Desjardins et al., 2020).

In the current study, mutations in the A-genome copy of *Ttzip4-A1* in tetraploid wheat produced a phenotype almost indistinguishable from wild type, and whilst there was a significant decrease in chiasma (representing crossover) frequency in *Ttzip4-B1* and *Ttzip4-B2* single, and *Ttzip4-A1B2* and *Ttzip4-B1B2* double mutants, these were relatively small differences, with only an average of 1-2 extra univalents observed per cell. However, in *Ttzip4-A1B1* double mutants (where *TtZIP4-B2* is the only *ZIP4* copy present), crossover was reduced by 76-78% and plants were sterile, similar to observations in Arabidopsis and rice.

This was also similar to the crossover reduction (∼85%) previously seen in tetraploid wheat *Ttmsh4ab* double mutants (Desjardins et al., 2020). In that study, cytological methods showed that the remaining ∼15% of crossovers were accounted for by the class II pathway. Our data suggests that the group 3 copies, *TtZIP4-A1* and *TtZIP4-B1*, control most of the homologous crossover occurring in tetraploid wheat. Group 3 copies of *TaZIP4* are also thought to be predominantly responsible for the promotion of homologous crossover in hexaploid wheat (Martín et al., 2021). Interestingly, the triple mutant, *Ttzip4-A1B1B2*, with loss of function in all three *ZIP4* copies, shows a more than 95% reduction in crossover frequency and is completely sterile. If *ZIP4* is required for more than 95% of crossovers in tetraploid wheat, it suggests that when all three *TtZIP4* copies are present, they have a stronger effect on crossover formation than was previously reported for *ZIP4* in Arabidopsis and rice. This additional effect on crossover frequency means that, in terms of stabilizing crossover in wheat, *ZIP4* is dosage-dependent, which is unusual when compared to other major genes controlling crossover. In hexaploid wheat, the ancestral homeologous *ZIP4* copies on 3A, 3B, and 3D are still present and expressed (Griffiths et al., 2006; Alabdullah et al., 2019), suggesting that increased *ZIP4* gene dosage may bias recombination toward homologs rather than homeologs (Desjardins et al., 2020).

### *ZIP4* may affect the class II crossover pathway in addition to the class I pathway

Previous studies have indicated that when *ZIP4* is disrupted, around 85% of crossovers are eliminated, consistent with loss of all crossovers in the class I pathway (Chelysheva et al., 2007). Our study showed that when all three copies of *ZIP4* were disrupted in tetraploid wheat, loss of crossover increased to more than 95%, suggesting that the tetraploid wheat copy *TtZIP4-B2* may have an additional effect on the class II crossover pathway. In most organisms, the class I and class II crossover pathways appear to be independent, but our results suggest that this may not be the case in wheat. Interestingly, interference between class I and class II crossovers has already been shown in wild type tomato, although the mechanism of interaction is unknown (Anderson et al., 2014).

*ZIP4-B2* on chromosome 5B originates from chromosome 3B. On wheat polyploidization, a trans-duplication event caused the *ZIP4* gene on 3B to duplicate and diverge, with the resulting new gene inserting into chromosome 5B, where it became responsible for maintaining fertility (International Wheat Genome Consortium, 2018). When this occurred, the *ZIP4-B2* copy on 5B may have acquired a novel function, to stabilize the class II crossover pathway in addition to its existing effect on the class I pathway. In tetraploid wheat, the fact that deleting the *ZIP4-B2* gene in addition to the *ZIP4-A1* and *B1* genes reduces crossover by over 95%, suggests that the duplication and divergence of the *ZIP4* gene on tetraploid wheat polyploidization was required to increase homologous crossover levels, and so was an important event in tetraploid as well as hexaploid wheat evolution.

### Chromosome synapsis is delayed in the absence of all three *TtZIP4* copies

Although the underlying mechanism is unknown, crossover formation is functionally linked to assembly of the SC between parental chromosomes. ZIP4 functions as a scaffold protein and may also act as a ‘molecular chaperone’, and as a hub for multiple physical interactions between other ZMM proteins involved in crossover and components of the chromosome axis (De Muyt et al., 2018; reviewed Pyatnitskaya 2019; Pyatnitskaya, 2022). Thus, ZIP4 may provide a direct physical link between crossover-designated recombination intermediates and SC assembly (De Muyt et al., 2018). Originally known as SPO22, ZIP4 is a large tetratricopeptide repeat (TPR) protein (Perry et al., 2005; Tsubouchi et al. 2006). The mammalian orthologue is TEX11. TPR motifs in ZIP4 mediate protein-protein interactions and facilitate assembly of multiprotein complexes (D’Andrea and Regan, 2003).

In *Sordaria macrospora*, ZIP4 associates with the chromosome axes during early meiosis, and together with other ZMM proteins, directly mediates installation of the central SC region, forming chromosome bridges that draw the chromosomes sufficiently close together to allow initiation of SC polymerization (Dubois et al., 2019; Pyatnitskaya, 2022). At the end of pachytene, once synapsis is complete, ZIP4 localizes to sites of crossover complexes, where recombination then takes place. In Sordaria, if *ZIP4* is disrupted, most chromosomes can only partially coalign or are unable to coalign at all (Dubois et al., 2019). In budding yeast, in the absence of *ZIP4*, the SC protein ZIP1 is unable to polymerize along chromosomes, thus preventing SC assembly (Tsubouchi et al., 2006). However, mutation in the Arabidopsis gene *AtZIP4* does not prevent synapsis, showing that the two functions of ZIP4 (i.e., class I crossover maturation and synapsis) can be uncoupled (Chelysheva et al., 2007).

In the current study, we found that, in the absence of the group 3 *TtZIP4* copies, as well as in the wild-type control, initiation of synapsis occurs as normal during the early stages of the telomere bouquet and is virtually complete after the bouquet has dispersed. Thus, *ZIP4* function of group 3 *TtZIP4* copies resembles that in Arabidopsis and rice, in that *zip4* mutants show a similar reduction in crossover levels, but with synapsis largely unchanged. However, in the absence of all three *TtZIP4* copies, some attempts to initiate synapsis do appear to take place, but in most meiocytes the polymerization process does not progress normally from telomere regions, and synapsis is delayed. Subsequently, after the telomere bouquet has dispersed, synapsis has clearly been compromised and is not completed. This is consistent with our previous studies in hexaploid wheat, in which we observed that, in the absence of *Ph1*, homologous chromosome synapsis progresses more slowly (Martín et al., 2017).

Previously, we have shown, using hexaploid wheat and wheat-rye hybrids, that only homologs can synapse during the telomere bouquet stage whether *Ph1* is present or not. Homeologs can synapse only later, mostly at late zygotene and pachytene after the telomere bouquet has dispersed, but will not do so if homologs have already synapsed (Martín et al., 2014, 2017). Thus, the delay in homologous synapsis in the absence of *Ph1* provides an opportunity for homeologs to synapse after the telomere bouquet has dispersed. In a previous study on SC spreads in tetraploid wheat, more multivalent associations were observed in the *ph1c* mutant than in the wild type (Martinez et al, 2001). Given such associations can only occur after the telomere bouquet has dispersed (Martín et al., 2017), this observation suggests that, in the *ph1c* mutant, more synapsis is occurring after the bouquet stage, and hence in this mutant synapsis is also delayed. The effect on synapsis observed in the absence of *TtZIP4-B2* in this study confirms that *ZIP4-B2* is also responsible for the effect on synapsis reported in the *ph1b* and *ph1c* mutant. Given that *ZIP4-B2* arose by duplication and divergence of the chromosome 3B copy, and that 3B copies have little activity during synapsis, *TtZIP4-B2* probably gained this additional function during polyploidisation. This may have been the meiotic adaptation that was required to promote homologous pairing and synapsis during the telomere bouquet stage, ensuring synapsis and crossover only occurs between true homologs, and thus preserving polyploid fertility.

### *ZIP4* may facilitate early synapsis by promoting synchronized elongation of homologs

In many species, including plants, clustering of telomeres during the early prophase I of meiosis, and organisation of chromosomes into a ‘bouquet’ arrangement, is thought to facilitate early stages of homologous chromosome pairing (Scherthan, 2001). In the nematode *Caenorhabditis elegans*, ZIP4 is not present, but the DNA-binding protein HIM-8 promotes synchronous elongation of chromosomes during meiotic prophase, which appears to enable homologous chromosomes to associate and align along their entire lengths prior to synapsis (Nabeshima et al., 2011). A similar synchronous elongation of chromosomes has also been reported in wheat (Prieto et al., 2004). In a hexaploid wheat line, in which a segment of rye had been substituted for 15% of one of the wheat chromosome arms, visualization of the rye segments showed that they elongated synchronously, immediately before formation of the telomere bouquet and their intimate pairing (Prieto et al., 2004). However, in the *ph1b* deletion mutant, in 64% of meiocytes the two rye segments were observed to have different conformations (i.e., elongation of the rye segments was not synchronized) during early meiosis, and a similar proportion were also incorrectly paired. This suggested that promotion of homolog pairing is related to a synchronised conformational state.

In the current study, we have observed that *TtZIP4-B2* promotes early synapsis, but it remains to be seen whether delayed synapsis in the triple mutant is due to lack of synchronization of chromosome axis elongation. However, if ZIP4 function in early meiosis is analogous to that of HIM-8 in *C. elegans*, this would explain the observation made by Prieto et al., (2004) that *Ph1* (*ZIP4-B2*) promotes synchronized elongation. Studies in Sordaria reveal that initial chromosome interactions involve ZIP4 foci on homologous chromosomes (Dubois et al., 2019), so one explanation for the ability of *TtZIP4-B2* to promote homologous pairing is that *TtZIP4-B2* synchronizes homolog elongation. This would ensure similar homolog conformation, facilitating rapid association of ZIP4 foci and formation of pairing bridges, thus reducing the chance of homeologous pairing, which only occurs later in meiosis (Dubois et al., 2019; Alabdullah et al, 2021).

In summary, *ZIP4-B2* in wheat promotes homologous crossover and suppresses homeologous crossover. *ZIP4-B2* also promotes early synapsis at the telomere regions during the telomere bouquet stage and promotes the progression of synapsis so that it is completed. Thus, *ZIP4-B2* accounts for the two major phenotypes reported for *Ph1*. The presence of *ZIP4-B2* in the wheat genome also results in a doubling of grain number (Alabdullah et al., 2021). These studies explain why the duplication and divergence of *ZIP4-B2* was so important for the stabilisation of wheat as a polyploid and reveal the enormous contribution this duplication event has made to agriculture and human nutrition.

### Future studies

Our previous studies on the 59.3Mb region deleted in the *ph1b* mutant revealed that this region (termed the *Ph1* locus) also affects centromere behaviour during meiosis (Martinez-Perez et al., 2001). However, the consequence of this centromere effect on maintenance of wheat genome stability during meiosis is uncertain, and needs to be addressed. We have recently carried out mutant analysis that links *ZIP4-B2* with segregation of achiasmatic (univalent) chromosomes and balanced gametes (unpublished), but further work is required. Control of genes such as *ZIP4* that affect chromosome synapsis and crossover can be extremely useful in improving the allelic diversity of elite wheat cultivars via the introgression of useful genes from wild relatives. For example, the *ZIP4* mutant lines previously generated in hexaploid wheat can now be exploited in breeding as an alternative to *ph1b* mutant lines (Rey et al., 2017; Martín et al., 2021). It is hoped that in future breeding programs involving tetraploid lines, the *Ttzip4* mutant lines developed in the current study could be used instead of the Cappelli *ph1c* mutant. Going forward, it will also be important to identify whether there are *ZIP4* copy variants in tetraploid wheat with increased temperature tolerance during meiosis, given the profound effects of *ZIP4* on wheat meiosis.

## Acknowledgements

We are grateful to Martha Clarke for technical support in developing the CRISPR mutant; Mark Youles of TSL SynBio for supplying L0 Golden Gate components; the Germplasm Resource Unit at the John Innes Centre (JIC) for providing seed and JIC Horticultural Services staff for maintenance of plant material.

## Funding

This work was supported by the UK Biological and Biotechnology Research Council (BBSRC) through a grant as part of the ‘Designing Future Wheat’ (DFW) Institute Strategic Programme (BB/P016855/1) and response mode grant (BB/R0077233/1). MD-R thanks the contract “Ayudas Juan de la Cierva-Formación (FJCI-2016-28296)” of the Spanish Ministry of Science, Innovation and Universities.

---

**Supplementary Figure 1**

Column chart showing genotypic effects on meiotic metaphase I chromosomes of Kronos wild type and Kr3161 *TtZIP4-A1B1B2* control plants. The numbers of univalents, rod and ring bivalents and single and double chiasmata are shown. Multivalents are not shown. Error bars show standard error.

**Supplementary Table 1.**
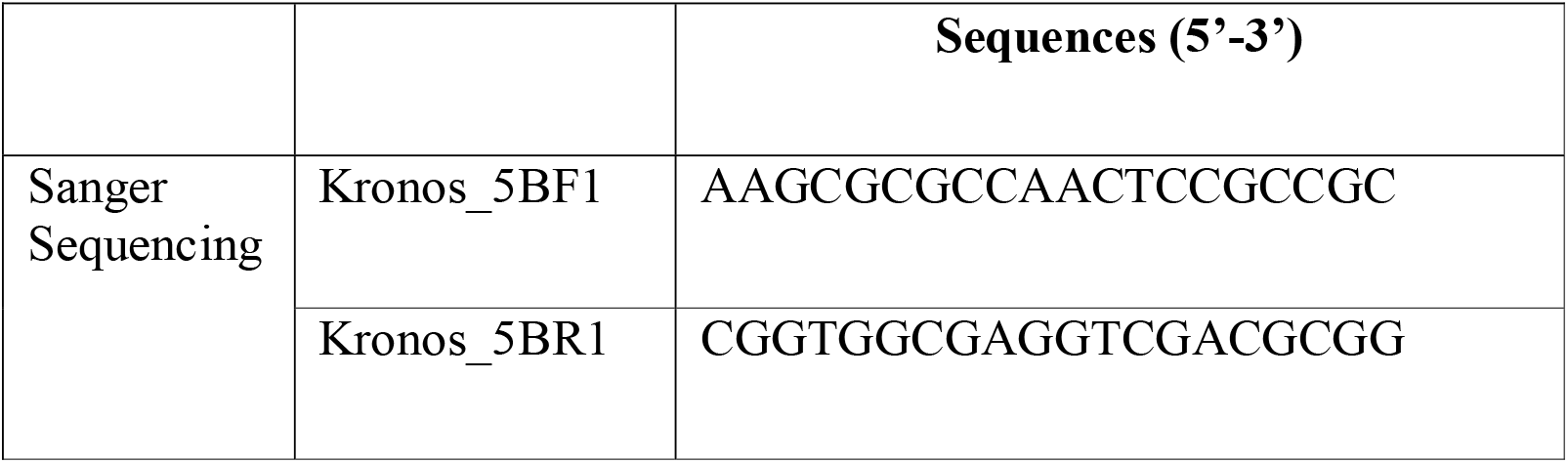
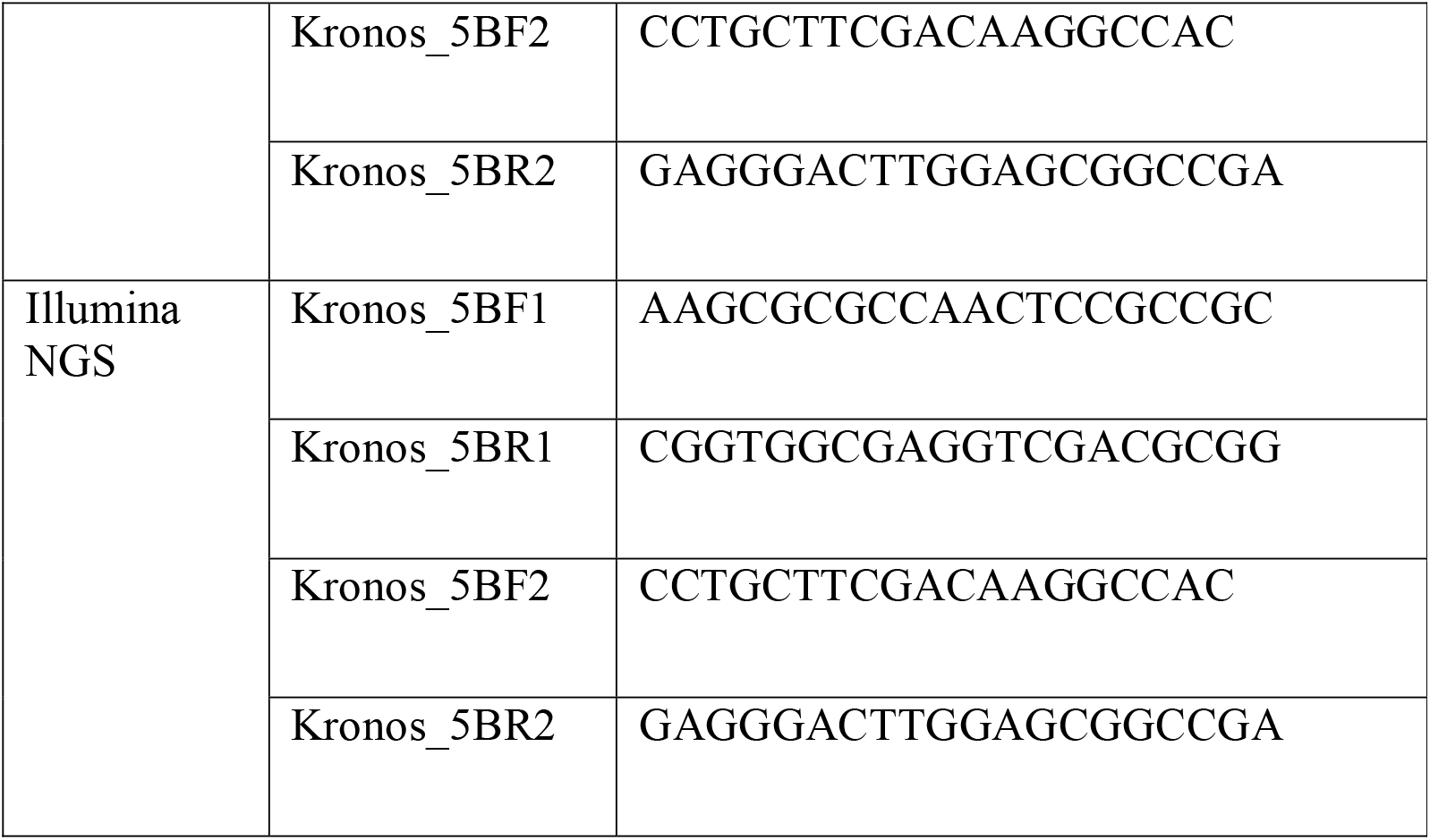
Primers for detection of CRISPR mutations in the *TtZIP4-B2* gene in the primary transgenics (T_0_) and subsequent generation (T_1_) using Sanger and Illumina sequencing

**Supplementary Table 2.**
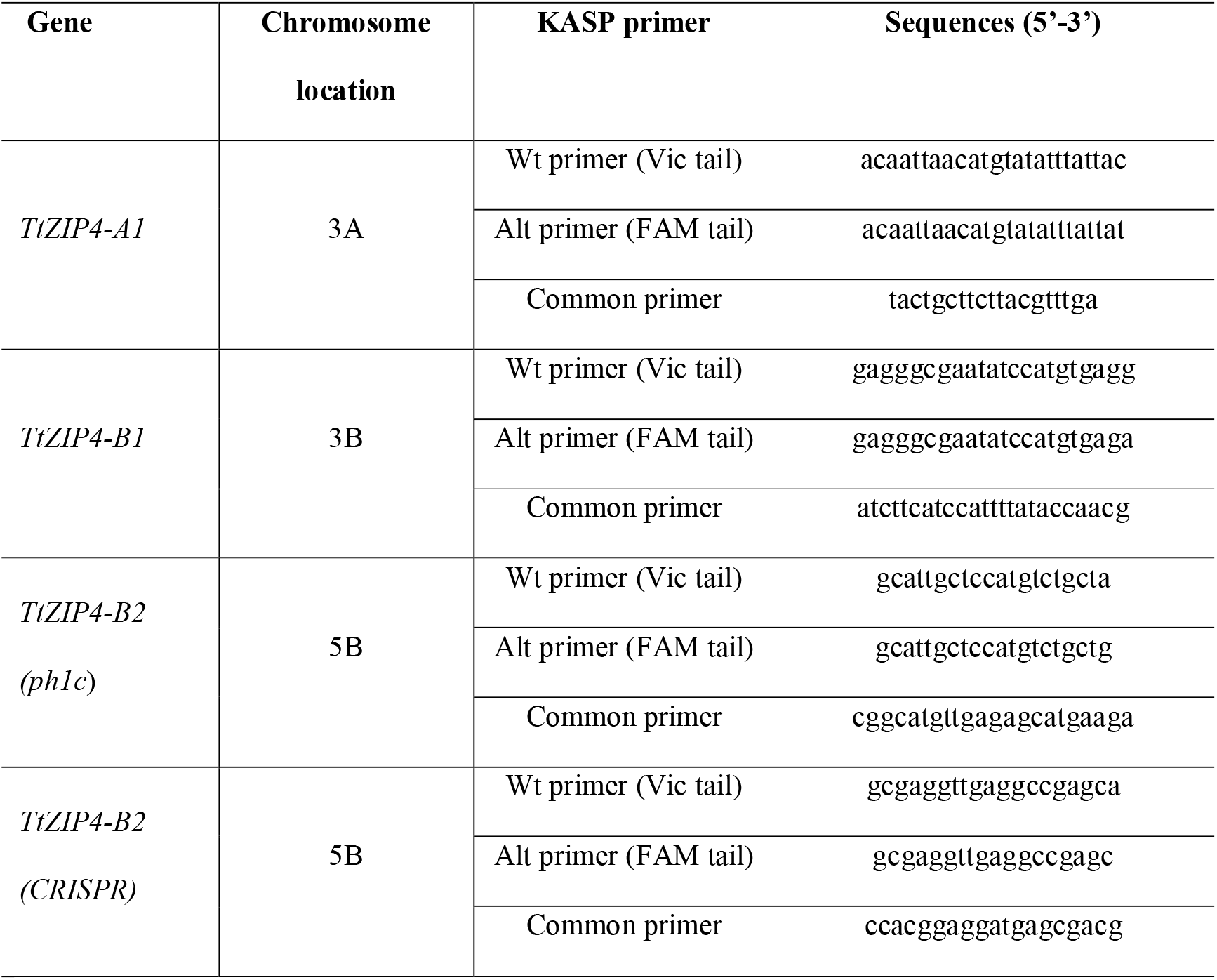
Genome-specific primer sequences for KASP genotyping *ZIP4* tetraploid wheat lines

**Supplementary Table 3.**
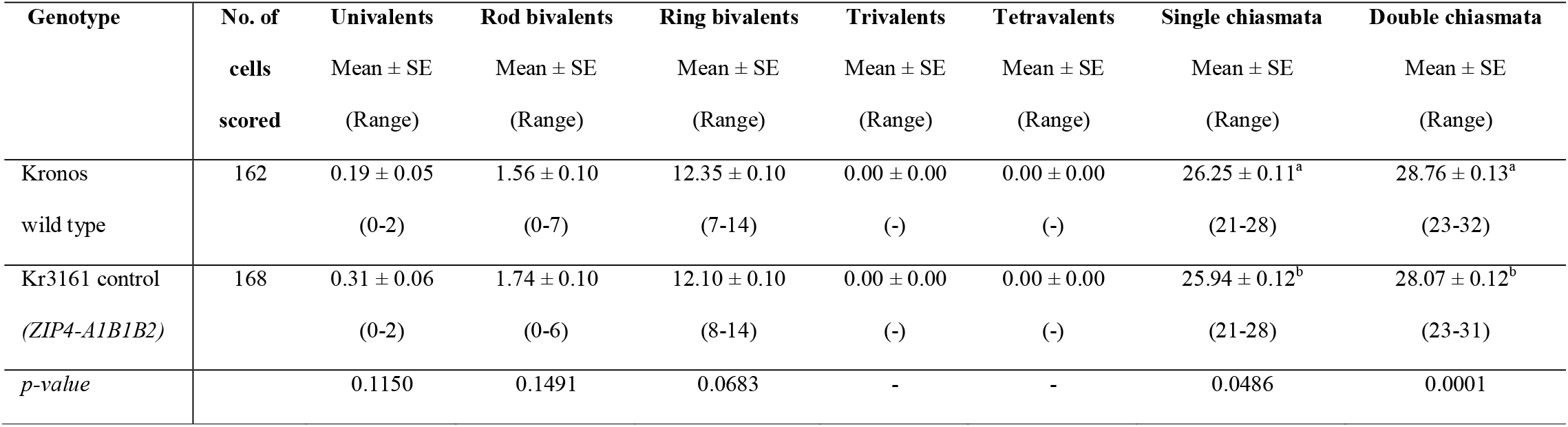
Genotypic effects on meiotic metaphase I chromosomes of Kronos wild type and Kr3161 *TtZIP4-A1B1B2* control plants. Mean numbers of univalents, rod and ring bivalents, trivalents and tetravalents were scored along with chiasma frequency scored as single and double chiasmata. Standard error (SE) values are shown. The range is shown in brackets. P-values < 0.05 indicate significant differences. Superscript letters a and b indicate where the significant differences lie. For scores with the same letter, the difference between the means is not statistically significant. If the scores have different letters, they are significantly different.

## Notes

### Competing Interest Statement

The authors have declared no competing interest.

