## Supplementary Figure 1 for "*ZIP4* is required for normal progression of synapsis and for over 95% of crossovers in wheat meiosis"

### Slide 1
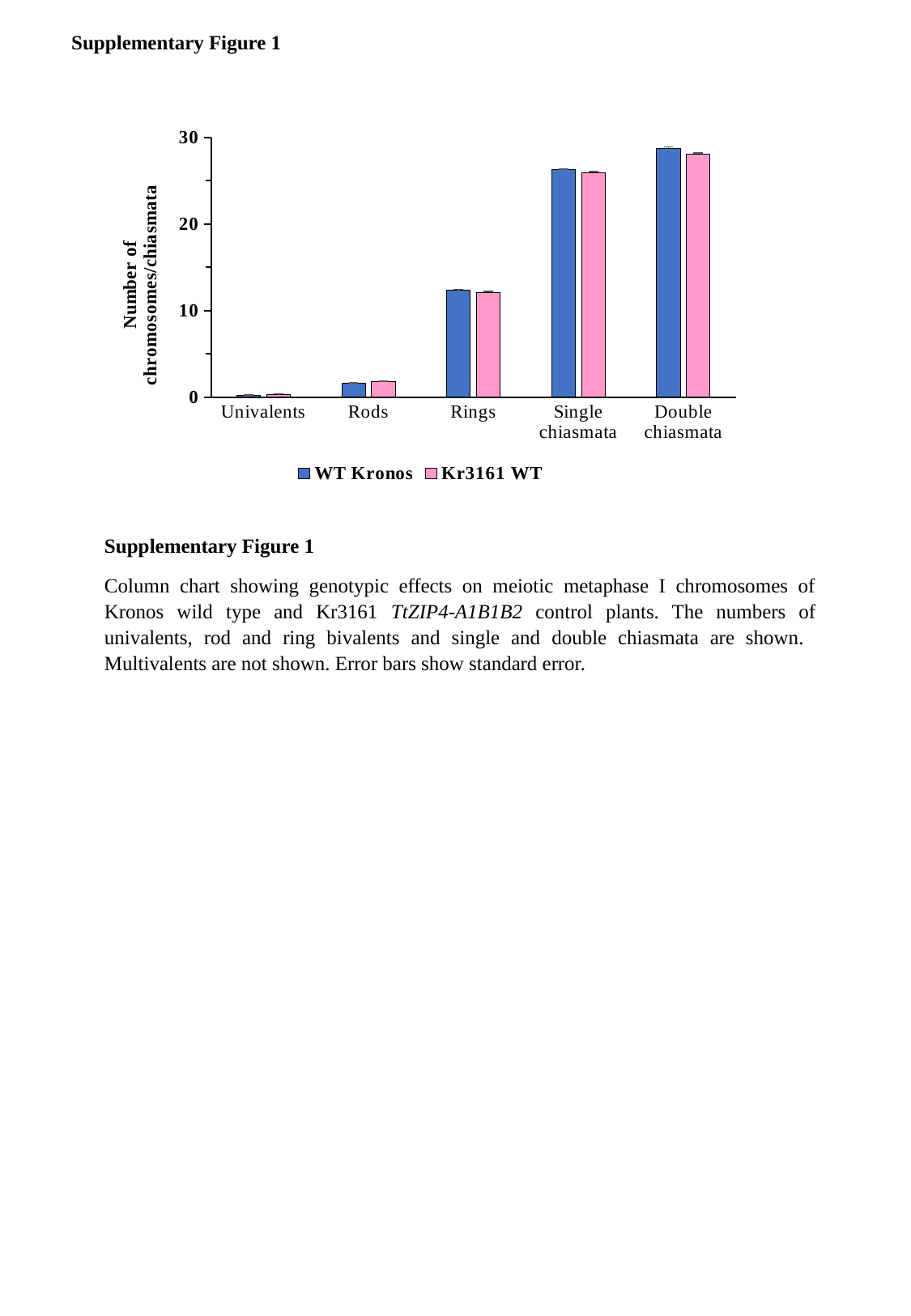

Supplementary Figure 1
#### Chart
| Category | WT Kronos | Kr3161 WT |
|---|---|---|
| Univalents | 0.19 | 0.31 |
| Rods | 1.56 | 1.74 |
| Rings | 12.35 | 12.1 |
| Single chiasmata | 26.25 | 25.94 |
| Double chiasmata | 28.76 | 28.07 |Supplementary Figure 1
Column chart showing genotypic effects on meiotic metaphase I chromosomes of Kronos wild type and Kr3161 TtZIP4-A1B1B2 control plants. The numbers of univalents, rod and ring bivalents and single and double chiasmata are shown. Multivalents are not shown. Error bars show standard error.
